# The mechanism of cell cycle dependent proteasome-mediated CdvB degradation in *Sulfolobus acidocaldarius*

**DOI:** 10.1101/2025.08.04.668399

**Authors:** Yin-Wei Kuo, Jovan Traparić, Sherman Foo, Buzz Baum

## Abstract

Protein degradation helps order events in the cell division cycle in eukaryotes, bacteria and archaea. This process is best understood in eukaryotes, where chromosome segregation and mitotic exit are triggered by APC/C and ubiquitin-regulated proteasome-dependent degradation of Securin and Cyclin B, respectively. Recent findings show that the archaeal proteasome also targets cellular substrates, including CdvB, for degradation in a cell cycle-dependent manner in *Sulfolobus acidocaldarius* - one of the closest experimentally tractable archaeal relatives of eukaryotes. Here, using CdvB as a model target protein to explore the mechanism of cyclic protein degradation, we identify the C-terminal broken winged helix of CdvB, which was previously shown to bind CdvA, as a domain that is sufficient to render a fusion protein unstable as cells transit from division phase to G1 phase. In parallel, we show that the rate of CdvB degradation accelerates during division, in part due to a cell cycle-dependent increase in the expression of the proteasome-activating nucleotidase (PAN), under the control of a cyclically expressed novel transcription factor, “CCTF1” (saci_0800), that can repress PAN expression. Taken together, our findings reveal the mechanisms by which archaea, despite lacking CDK/cyclin or APC/C proteins, regulate proteasome-mediated degradation to order events during cell division.

## Introduction

Orderly progression through a round of the cell division cycle requires certain proteins to be expressed and degraded at specific times (Fry & Yamano, 2006; Fatima *et al*, 2022; Tarrason Risa *et al*, 2020). In eukaryotes, a wave of protein degradation at mitotic exit resets the cell cycle and triggers chromosome segregation, cytokinesis, and replication origin licensing. This process is initiated upon satisfaction of the spindle assembly checkpoint via the recruitment of Cdc20 to the CDK-activated Anaphase Promoting Complex (APC/C) (Musacchio, 2015). The Cdc20-APC/C complex then ubiquitylates proteins that contain a destruction-box motif (RXXLXXXXN), including CyclinA, CyclinB, and Securin, targeting them for proteasome-mediated degradation (Glotzer *et al*, 1991; McAinsh & Kops, 2023). As cells exit mitosis and enter G1, Cdc20 is then swapped out for Cdh1, which targets additional substrates of the APC/C that typically contain a KENXXXN box motif, like Cdc20, for ubiquitylation and degradation (Pfleger & Kirschner, 2000; Davey & Morgan, 2016). This extended wave of cyclic proteasome-mediated protein degradation is therefore achieved by: (i) regulation at the level of a substrate’s affinity for the degradation targeting machinery ensuring, for example, that the same complex ubiquitylates and degrades Cyclin A before Cyclin B, and (ii) at the level of the degradation machinery itself, with Cdh1-APC/C taking over from Cdc20-APC/C at late stages of mitotic exit.

Despite lacking eukaryotic-like cell cycle regulators such as cyclins, CDKs or the APC/C, Thermoprotei like Sulfolobales have an ordered cell cycle that appears superficially similar to that of many eukaryotes (Cezanne *et al*, 2024; Lindås & Bernander, 2013). While it is not yet known how this is regulated, progression from G2 into G1 in *Sulfolobus* is accompanied by a wave of proteasome-mediated protein degradation that resembles the one observed in eukaryotes exiting mitosis and undergoing cytokinesis. A key target of the *Sulfolobus* proteasome at division is CdvB (Tarrason Risa *et al*, 2020; Liu *et al*, 2025), an ESCRT-III homolog (Samson *et al*, 2008; Lindås *et al*, 2008). The timely degradation of CdvB is critical for orderly cell cycle progression, since CdvB must first accumulate to form a medial polymeric ESCRT-III ring before DNA segregation and cytokinesis can occur (Parham *et al*, 2025). Then, a few minutes later, CdvB polymer disassembly and concomitant CdvB degradation are required for constriction of the cytokinetic CdvB1/B2 ring and abscission (Tarrason Risa *et al*, 2020; Harker-Kirschneck *et al*, 2022).

The entire sequence of events that leads to cell division in *Sulfolobus acidocaldarius* are as follows: As G2 cells prepare for division they begin to express a wave of cell division proteins, which include CdvA, CdvB, CdvB1 and CdvB2, and Vps4. CdvA is the first to accumulate (Samson *et al*, 2011). CdvA forms a medial ring that defines the division plane (Samson *et al*, 2011; Lindås *et al*, 2008). CdvA then recruits CdvB, through the interaction of the CdvB broken winged helix domain with the E3B region of CdvA (Samson *et al*, 2011). Subsequently, CdvB1 and CdvB2 are recruited to the ESCRT-III ring, leading to the formation of a precisely positioned composite division ring (Hurtig *et al*, 2023). This is coupled to DNA segregation (Parham *et al*, 2025). Once cells are ready to divide, the CdvB polymer is then disassembled by the AAA-ATPase Vps4, and the monomeric CdvB released is then degraded by the proteasome via a process that requires the proteasome-activating nucleotidase (PAN) (Hurtig *et al*, 2023; Tarrason Risa *et al*, 2020), an archaeal homolog of the 19S proteasome subunit (Maupin-Furlow, 2011; Forouzan *et al*, 2012). The loss of CdvB then allows the CdvB1/B2 cytokinetic ring to constrict (Harker-Kirschneck *et al*, 2022).

Although the formation and the timely destabilization of the CdvB ring are both known to play critical roles during division in *S. acidocaldarius*, it is not yet clear how cells ensure that CdvB is targeted for proteasome-mediated degradation at precisely the right time. Here, in exploring the mechanism of cyclic CdvB regulation in *S. acidocaldarius*, we map the regions of CdvB that contribute to its cyclic instability. This identifies a portion of the C-terminus of CdvB that renders the protein unstable across the cell cycle and identifies the broken winged helix domain of the protein, which was previously shown to bind CdvA, as playing a role in selectively stabilizing the protein during the pre-division phase of division ring assembly. In addition, we demonstrate a role for transcription factor-mediated changes in the levels of the PAN that contributes to the increase in CdvB degradation as cells initiate division. Taken together, these data reveal how sequences in the tail of CdvB that target the protein for degradation combine with general changes in the levels of the protein degradation machinery to regulate cell cycle progression in *Sulfolobus*.

## Results

### The C-terminal region of CdvB contributes to its rapid degradation

To understand the molecular origin of the rapid proteasome-mediated degradation of CdvB as *Sulfolobus* cells switch from a phase of CdvB ring assembly to CdvB1/B2-dependent ring constriction, we began by using flow cytometry to compare the degradation kinetics of CdvB relative to its two homologs, CdvB1 and CdvB2 (Fig. 1). This revealed, as previously described (Hurtig *et al*, 2023), that while the three proteins form a composite ring of homopolymers in cells preparing to divide, the entire cellular pool of CdvB is degraded before division is complete (Figure 1A). By contrast, CdvB1 and CdvB2 are gradually degraded several minutes later as cells pass from G1 into S phase (Fig. 1B, C). Interestingly, the G1/S degradation of CdvB1 and CdvB2 also depends on the proteasome since, like CdvB, the rate of their degradation in G1 could be reduced by the addition of bortezomib, a proteasome inhibitor, to the medium (Fig. S1). These data indicate that different proteins are targeted to the proteasome at different times in the *Sulfolobus* cell cycle – as is the case for substrates of the proteasome in eukaryotic cells (Morgan, 2007).

**Figure 1.**
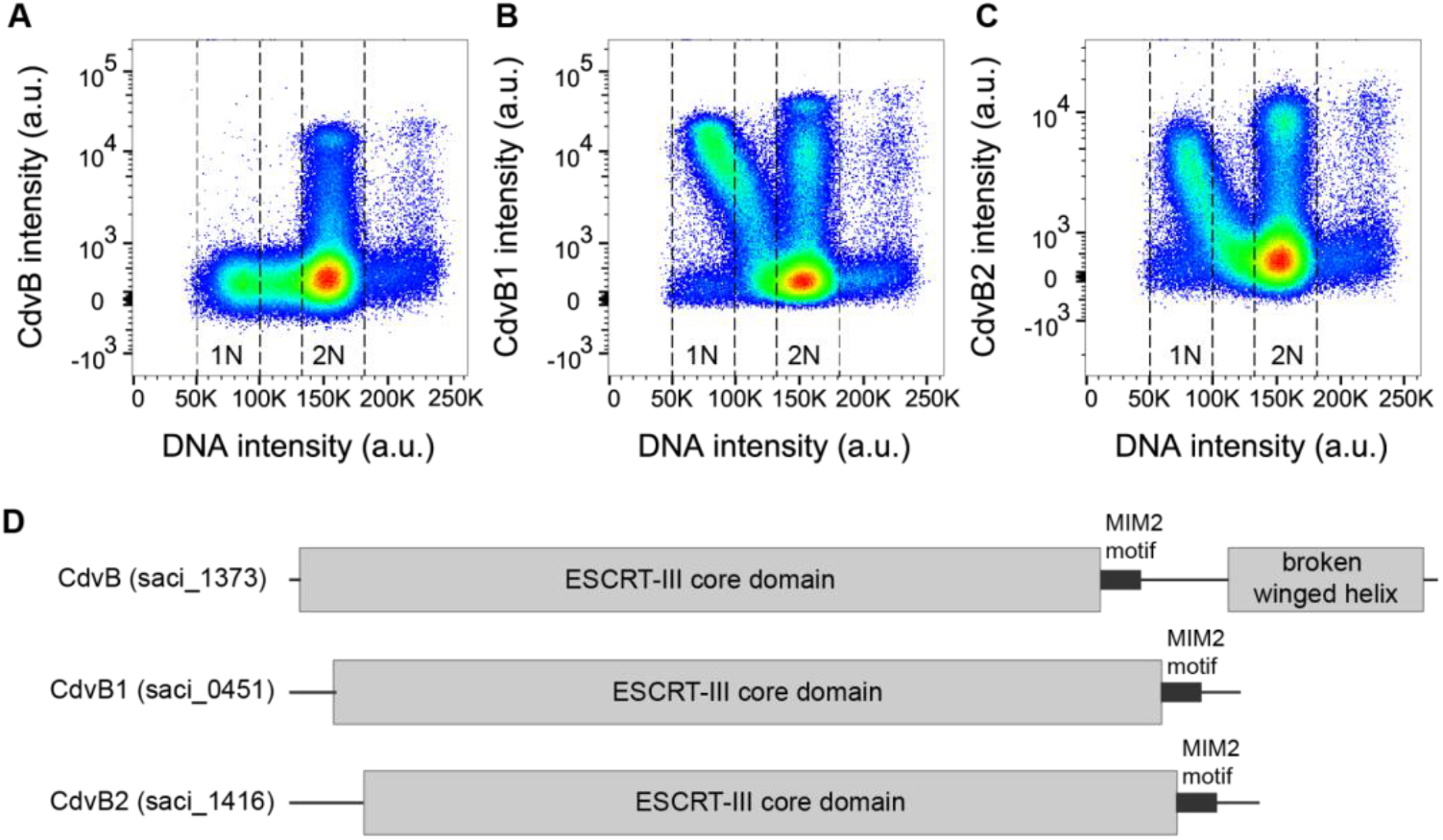
The ESCRT-III homologs of *Sulfolobus acidocaldarius*. **(A-C)** Representative flow cytometry analysis of asynchronous cultures of the uracil auxotrophic strain MW001 showing the levels of CdvB (A), CdvB1 (B), or CdvB2 (C) plotted against DNA content. Cells in G1 and G2 phases of the cell cycle are indicated as 1N and 2N respectively. Each dot represents a single event with the density gradient going from blue to red (n=2.5×10^5^ events each). **(D)** Schematic diagram of the architecture of the three ESCRT-III homologs showing the conserved ESCRT-III core domain, the MIM2 motif, the connecting linker, as well as the broken winged helix motif of CdvB.

These substrate-specific differences in the timing of proteasome-dependent protein degradation in dividing *Sulfolobus* cells prompted us to compare the domain architecture of the three homologs (Fig. 1D). Inspection of the sequences showed that CdvB has a longer C-terminal domain that contains a MIM2 motif, a linker region and an additional domain containing a broken winged helix (Fig. S2, Fig 1D), which is conserved across Sulfolobales (Makarova *et al*, 2024) and has been reported to interact with the non-ESCRT-III division protein, CdvA (Samson *et al*, 2011).

To test whether the C-terminal region of CdvB contains information that is sufficient to induce fast proteasome-mediated degradation during division, we fused the C-terminal region of CdvB to an N-terminally HA-tagged LacS protein: a beta-galactosidase variant taken from *Saccharolobus solfataricus* (see Fig. S3 for details on the choice of LacS). We then expressed this fusion protein on a plasmid containing the arabinose-inducible promoter and, following induction with 0.2% arabinose, used Western blotting to compare the expression of the HA-LacS-CdvB^C-term^ fusion protein with LacS fused to the C-terminal tail of CdvB1 and CdvB2, or an HA-LacS control.

While the LacS and the LacS fusion proteins carrying CdvB1 or CdvB2 C-terminal regions accumulated to high levels in asynchronous populations of cells, the CdvB C-terminal tail fusion was expressed at much lower levels under the same conditions (Fig. 2A). This implies that the C-terminus of CdvB contains information that renders a fusion protein unstable. We confirmed that this was due to proteasome-mediated protein degradation by showing that the HA-LacS-CdvB^C-term^ fusion protein rapidly accumulated following the addition of a proteasome inhibitor, bortezomib (Fig. S4).

**Figure 2.**
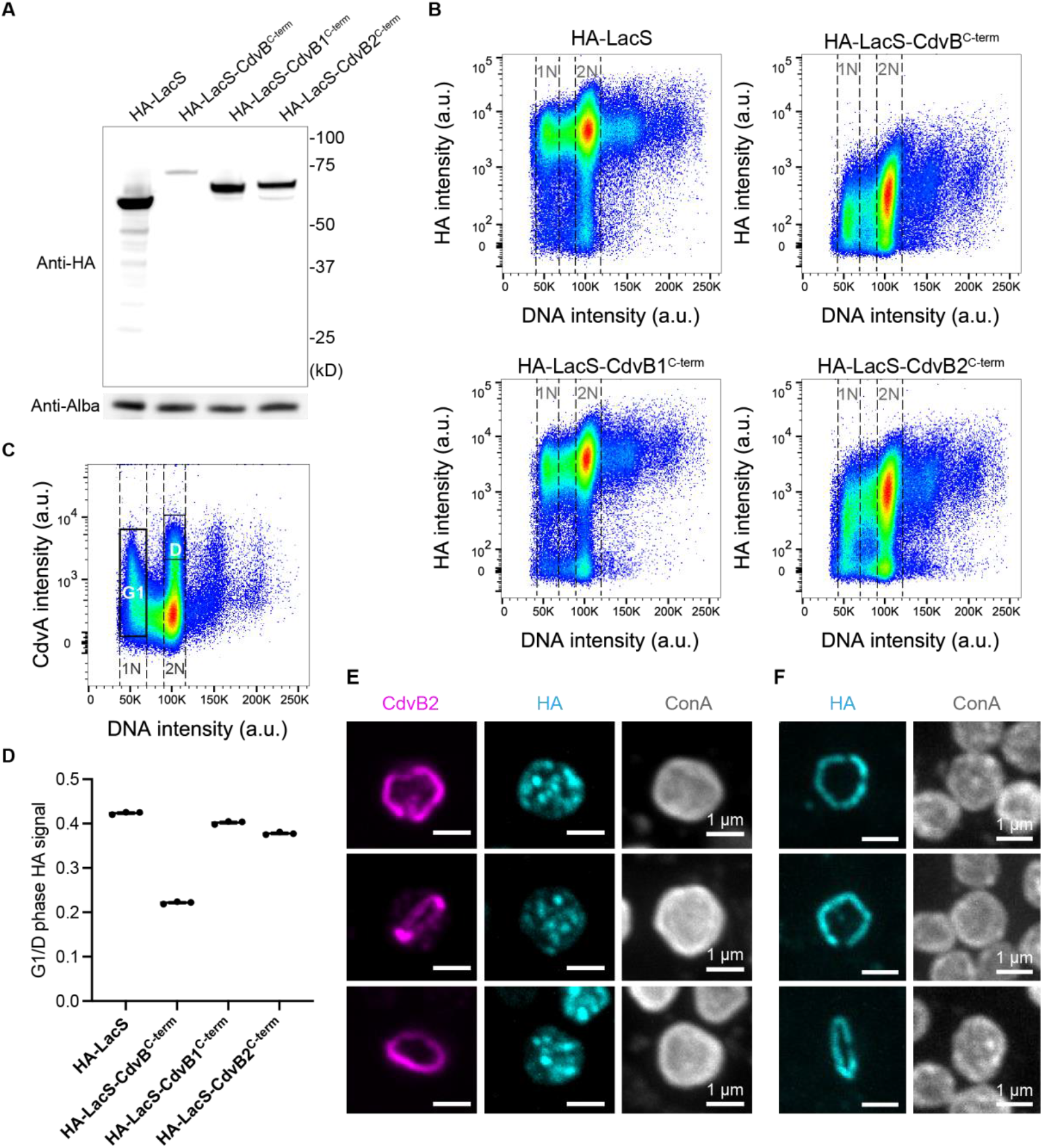
Effects of the C-terminal region of CdvB on LacS stability. **(A)** Representative Western blot of HA-tagged LacS fused with the C-terminal regions of the ESCRT-III proteins CdvB, CdvB1 or CdvB2 under the expression of the arabinose promoter. Alba was used as the loading control. **(B)** Flow cytometric analysis of HA-tagged LacS fused with the C-terminal regions of the ESCRT-III proteins CdvB, CdvB1 or CdvB2, plotted against DNA content. Cells in G1 and G2 phases of the cell cycle are indicated as 1N and 2N respectively. Each dot represents a single event with the density gradient going from blue to red (n=2.5×10^5^ events each). **(C)** Representative analysis of the CdvA levels in asynchronous *S. acidocaldarius* cells, showing the division phase (D-phase) and G1 phase. **(D)** Quantification of the HA signal from (B) as a ratio of the average level in G1 and D-phases. **(E, F)** Example immunofluorescence images of HA-LacS-CdvB^C-term^ (E) and full-length HA-CdvB overexpression cells (F) in the pre-constriction stage.

Next to assess the impact of the CdvB tail on cyclic protein degradation, we used flow cytometry to evaluate the levels and timing of HA-LacS-CdvB^C-term^ protein accumulation relative to HA-LacS-CdvB1/B2^C-term^ proteins. Consistent with the Western blotting analysis, this flow cytometric analysis revealed that the CdvB^C-term^ fusion protein had the lowest levels of HA expression (Fig. 2B). We then separated the population into G1 and Division phases (D-phase) using CdvA and DNA content as a guide (Fig. 2C) to obtain information about the levels of each fusion protein as they progress through the cell cycle. Interestingly, this analysis revealed that the HA-LacS-CdvB^C-term^ fusion protein is rapidly degraded during the transition from the D-phase (2N DNA content and high CdvA) into G1 phase transition (1N DNA content), resulting in an approximately 5-fold decrease in protein (equivalent to a G1/D phase HA-signal ratio of ∼20% (Figure 2D)). By contrast, the G1/D ratio for LacS, or the CdvB1^C-term^ or CdvB2^C-term^ fusion proteins was closer to 50% - the value expected for a stable protein that is evenly partitioned between two daughter cells during division (Fig. 2D). These data show that the C-terminal region of CdvB contains sufficient information to confer cyclic proteasome-dependent protein degradation on a fusion protein.

For endogenous CdvB to be degraded prior to division, the protein must be first removed from the polymeric ESCRT-III ring by the activity of the AAA-ATPase Vps4, as cells switch from a ring assembly phase to a ring constriction phase (Hurtig *et al*, 2023; Liu *et al*, 2025). To determine whether a similar process might contribute to the degradation of the HA-LacS-CdvB^C-term^ fusion protein, we used fluorescent microscopy to test if the C-terminal tail of CdvB is sufficient to target LacS to the medial ESCRT-III ring in dividing *Sulfolobus* cells. This was not the case. HA-LacS-CdvB^C-term^ appeared to be cytoplasmic and was not recruited to the division ring - consistent with it lacking the ESCRT-III domain required for polymerization (Fig. 2E), while overexpressed full length CdvB-HA can be recruited to the division ring (Fig. 2F). This indicates that the degradation of the fusion protein carrying the CdvB C-terminus is independent of ESCRT-III polymerization dynamics and must be regulated in some other way as cells initiate cell division.

### Dissecting the region of CdvB required for cyclic degradation

To map the specific signal(s) at the C-terminal region required for the cyclic degradation of LacS-CdvB^C-term^, we generated a series of truncation constructs removing the α5 helix of the ESCRT-III core domain, the MIT-interacting motif (MIM2), and the broken winged helix domain in different combinations. We then ran a flow cytometric analysis (Fig. 3, Fig. S5) to compare the levels of different fusion proteins in dividing cells (D-phase) and in G1 cells – using the G1/D-phase HA-signal ratio as a quantitative read-out of cyclic degradation.

**Figure 3.**
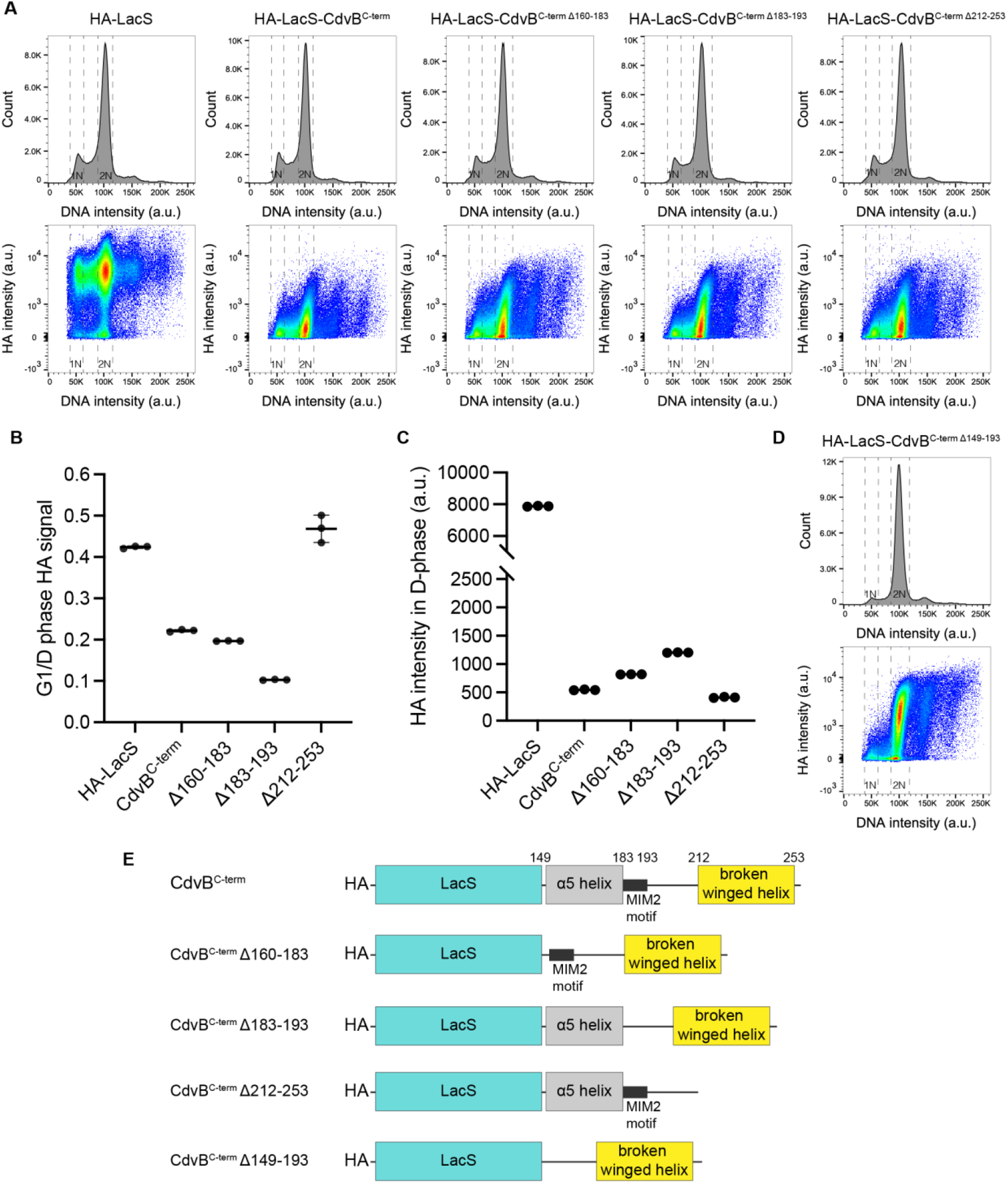
Identification of the region required for cyclic degradation at the C-terminal region of CdvB. **(A)** Representative flow cytometry histograms and scatter plots of HA-LacS fused with the CdvB^C-term^ truncation constructs. **(B)** Quantification of cyclic degradation of indicated truncation constructs using the HA signal ratio in G1/D-phase. **(C)** Average background subtracted HA intensities of the indicated constructs in D-phase. **(D)** Representative flow cytometry histogram and scatter plot of the Δ149-193 construct. **(E)** Schematic diagram of the LacS-CdvB^C-term^ constructs used in this study.

As expected, the inclusion or removal of α5 helix had little, if any, effect on the degradation of the C-terminal fusion construct since this is part of the ESCRT-III core domain (Fig. 3A, B, Fig. S5A, C). By contrast, the instability that is characteristic of the CdvB tail was lost entirely in the Δ194-261 deletion construct, which retains the α5 helix and MIM2 domain but lacks everything else, and in the 194-211 construct, which lacks everything but the linker connecting the MIM2 domain and the broken winged helix (Fig. S5). This identifies regions in the C-terminal domain of CdvB that render the protein unstable across the cycle.

When we focused our analysis on the broken winged helix domain, however, we observed a distinct effect of this region of the C-terminal tail on cyclic protein stability. While a fusion protein that lacked the broken winged helix motif but retained the MIM2 and linker regions still proved unstable (Δ212-253), it failed to undergo cell cycle stage-specific degradation during the transition from D-phase to G1 phase (Fig. 3A, B). As a result, levels of the (Δ212-253) fusion protein were only reduced by ∼50% upon passage from D-phase into G1, like the LacS and CdvB1 and CdvB2 fusion protein controls (Fig. 3B). Furthermore, when we analysed the data in more detail it became clear that this was due to the ability of the broken winged helix domain to increase fusion protein stability in D-phase of the cycle (Fig. 3C, Fig. S6). Somewhat paradoxically, this implies that the cyclic degradation of fusion proteins carrying the C-terminal tail of CdvB is due to the ability of the broken winged helix to stabilize proteins during early division, whilst it cannot do so at later stages. Since the broken winged helix domain is expected to interact with CdvA (Samson *et al*, 2011), it is possible that the destabilization of CdvB as cells pass from G2 into G1 in part reflects the loss of binding of the CdvB C-terminal tail to CdvA during the transition from the ring assembly to the ring constriction phase of division.

Through this analysis, we also noted that levels of the fusion protein in D-phase were even higher in cells carrying Δ149-193 deletion constructs that retain the broken winged helix domain, but which lack the α5-helix and MIM2 domain (Fig. 3D; Fig. S6 bottom). In this case, however, the G1 population was markedly reduced, implying a block in cell division (Fig. 3D). This is striking, as it mirrors the effects of expressing a CdvA construct that lacks the CdvB interaction site (Parham *et al*, 2025), which inhibits the transition from pre-division to constriction. This implies that the expression of a fusion protein that carries the broken winged helix domain that can bind CdvA at very high levels may interfere with the binding of endogenous CdvB with CdvA during the ring assembly phase.

Taken together, these results suggest that the cyclic degradation of CdvB during division is regulated by sequences that destabilize the protein across the cycle, together with a broken winged helix domain that preferentially stabilizes the protein during the ring assembly phase of division.

### The cyclic degradation of CdvB accords with cyclic expression of proteasome-activating nucleotidase (PAN)

After identifying the broken winged helix as part of the molecular signal that regulates cyclic degradation of CdvB, we next examined if there are changes in the protein degradation machinery itself during the transition from ESCRT-III ring assembly to cytokinesis. We began by investigating the involvement of the proteasome-activating nucleotidase (PAN) in the degradation of CdvB, since previously published work had implicated it in the proteasome-mediated degradation of CdvB (Tarrason Risa *et al*, 2020). Consistent with this earlier work, overexpression of the ATP-hydrolysis deficient mutant (E237Q) of PAN led to the increased division failure as manifested by the accumulation of polyploid cells (Fig. 4A, B). We also observed a residual pool of CdvB protein in newly divided G1 cells expressing this dominant negative PAN, which was absent from G1 cells in the empty vector control strain (Fig. 4A black boxes, 4C) and from G1 cells in the MW001 background strain (Tarrason Risa *et al*, 2020). While we did not previously observe G1 cells with residual CdvB after treatment with high dosage of the proteasome inhibitor (Fig. S7A-C), which blocks cell division (Tarrason Risa *et al*, 2020), a population of G1 cells containing residual CdvB was observed when cells were exposed to low dosage of bortezomib (Fig. S7B). These data imply that CdvB accumulates in G1 cells if its degradation is partially compromised.

**Figure 4.**
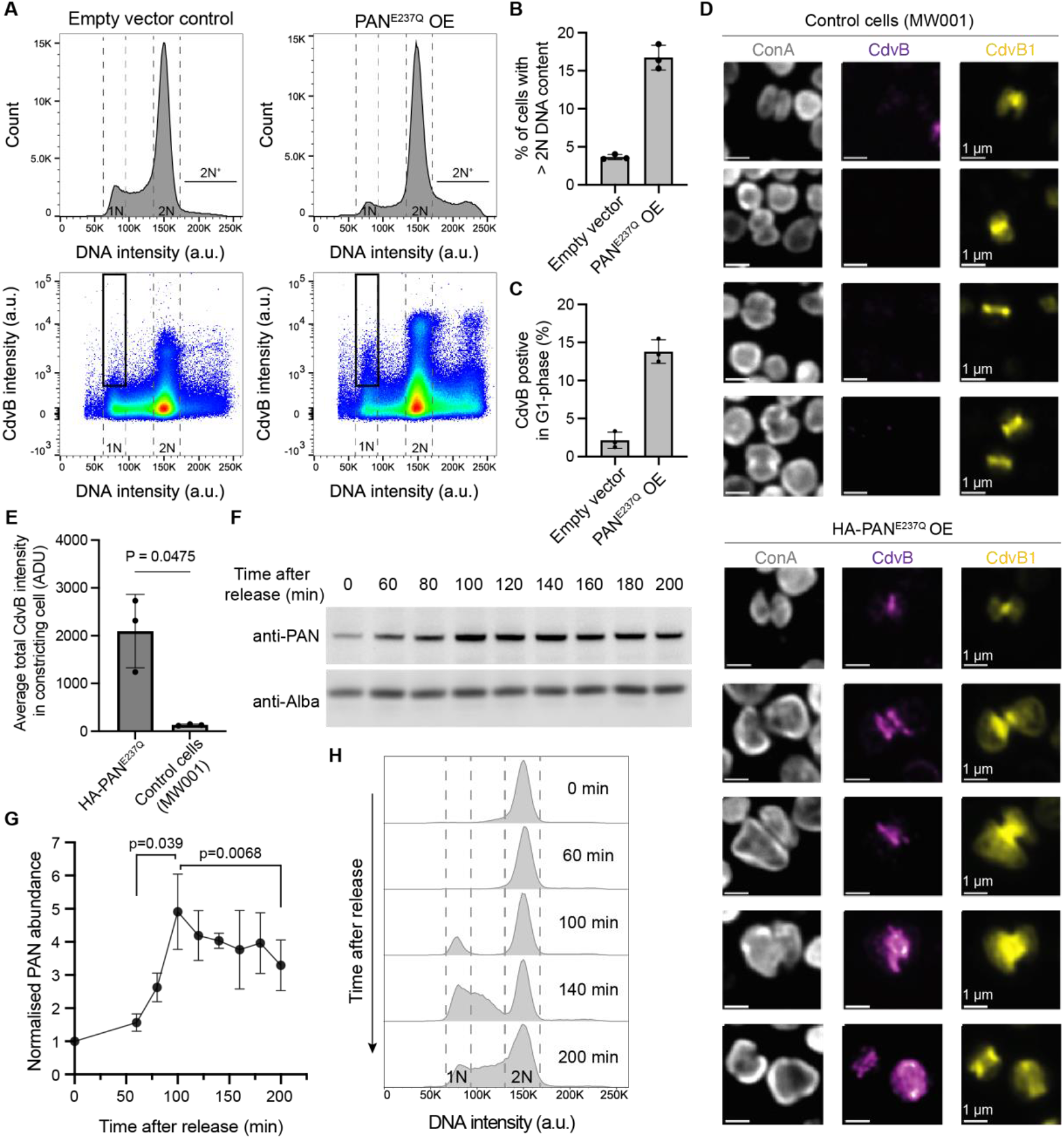
Disruption of PAN activity decreases degradation of CdvB. **(A)** Representative flow cytometry histograms and scatter plots of the dominant negative mutant of PAN (PAN^E237Q^) overexpression after 4 hr addition of arabinose; n=5.0×10^5^ events each. **(B, C)** Percentage of cells with more than 2N DNA content (B) and CdvB positive cells in G1 phase (C) from flow cytometry analysis in (A). **(D)** Example immunofluorescence images (maximum projection) of membrane constricting cells from MW001 background strain control (top) and HA-PAN^E237Q^ overexpression (bottom). **(E)** Quantification of total CdvB intensity of constricting cells in (D). Welch t-test; N=3 biological replicates. ADU: analog digital unit. **(F-H)** PAN shows cyclic expression in synchronized cells (MW001) after release from acetic acid induced cell cycle arrest with representative Western blots showing the level of PAN and Alba loading control (F). The corresponding flow cytometry histograms of DNA content are shown in (H). Quantification of PAN expression level from the Western blots is shown in (G). The relative PAN protein level is normalised by loading control (Alba) and time point = 0 min. Error bars: means ± SDs; ratio paired t-test, N=3 biological replicates.

To determine how the loss of CdvB degradation influences division, we fixed, stained and imaged cells by immunofluorescent microscopy. This revealed that the expression of the PAN^E237Q^ mutant prevented the degradation of CdvB that typically precedes membrane constriction in *Sulfolobus* cells (Fig. 4D, E). In many cases, this appeared to lead to a failed division, resulting in an increase in cell size (Fig. 4D) and the accumulation of polyploid cells (Fig. 4B).

Encouraged by these results implicating PAN in the timely degradation of CdvB, we performed a cell cycle synchronization experiment to determine whether PAN activity might change during cell cycle progression in *Sulfolobus* in a way that contributes to the change in CdvB stability. When cells were released from an acetic acid arrest and followed as they entered the cell cycle, we observed an increase in the levels of PAN by Western blotting (Fig. 4F). Its expression peaked at ∼100 min after release, coinciding with D-phase (Fig. 4F-H), indicating that PAN is subject to cell cycle-dependent expression.

Concomitant with the current study, we discovered that a putative ArsR family transcription factor, *saci_0800*, that when over-expressed from the arabinose promoter to induce flat expression caused a phenotype similar to the PAN^E237Q^ dominant negative, in which G1 cells accumulated CdvB protein (Fig. 5A-C). We named this factor ‘Cell cycle Control Transcription Factor 1’ (CCTF1). The CCTF1 gain of function phenotype led us to consider whether it might influence PAN expression, especially since transcript levels of CdvB were slightly decreased in the CCTF1 overexpression strain (Fig. 5D). This prediction was borne out by Western blotting and RT-qPCR analyses, which showed that CCTF1 overexpression strongly reduced the expression of PAN on both transcriptional and protein level (Fig. 5E, F). Consistent with this, we also observed CCTF1 overexpression preventing the complete degradation of CdvB during membrane constriction (Fig. 5G, H). Thus, overexpression of CCTF1 downregulates the expression of PAN and consequently decreases the degradation activity against CdvB during D-phase.

**Figure 5.**
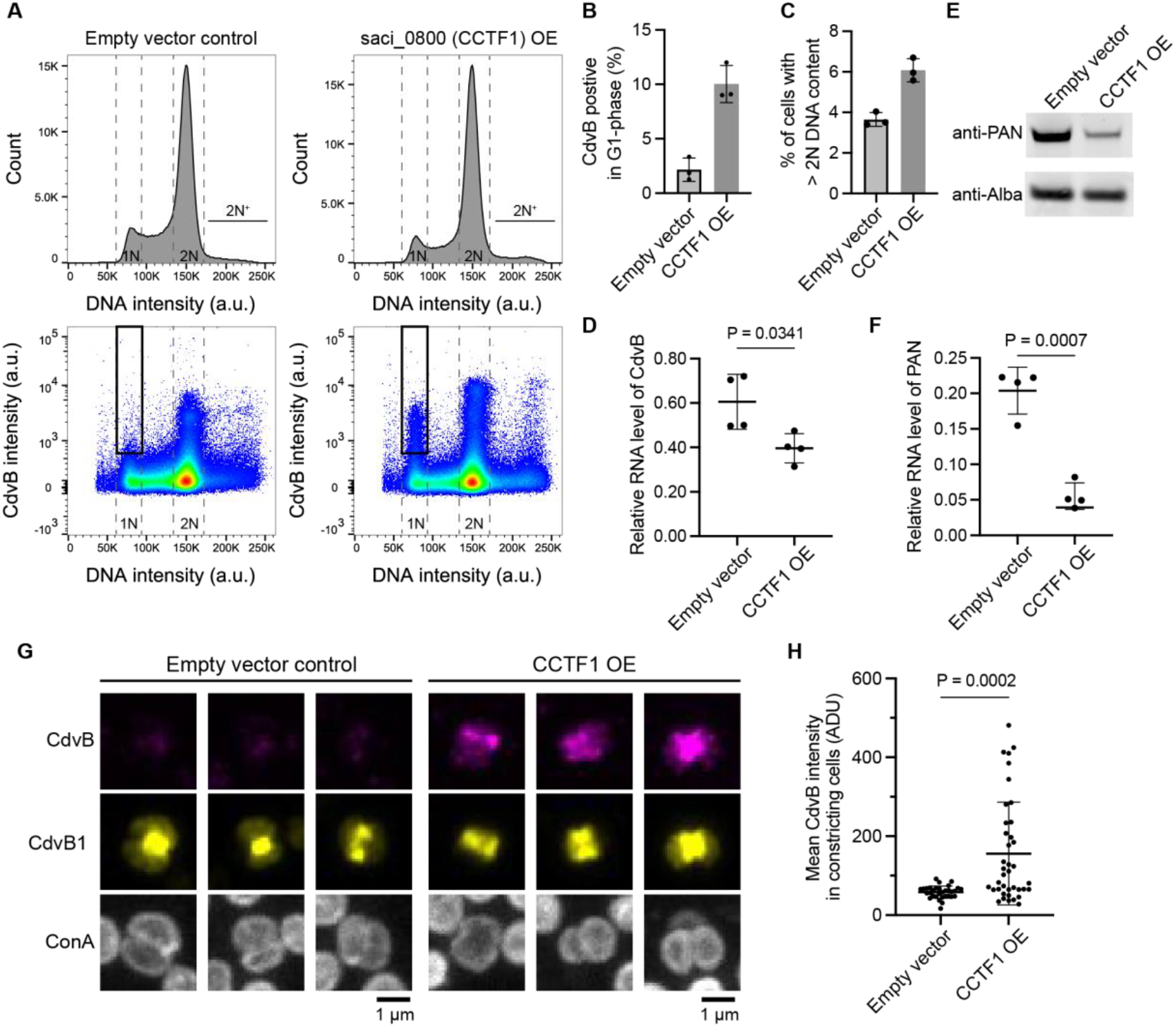
Overexpression of CCTF1 suppressed degradation of CdvB. **(A-C)** Example flow cytometry histograms and scatter plots of CCTF1 (saci_0800) overexpression (A) with quantification of CdvB positive cells in G1 phase (B) and percentage of cells with more than 2N DNA content (C). Note that the empty vector controls here are the same dataset as in Fig. 4A-C and are repeated here for clarity. **(D)** Representative Western blots of PAN protein signal in empty vector control and CCTF1 overexpression. **(E, F)** RT-qPCR analysis of CdvB (E) and PAN (F) in empty vector and CCTF1 overexpression strains after 4 hr addition of arabinose. Welch t-test, N=4; error bars: mean±SDs. **(G)** Example of the immunofluorescence maximum projection images of CCTF1 overexpression and empty vector controls in the cell constriction stage. **(H)** Quantification of the mean CdvB intensity in the membrane constricting stage by immunofluorescent imaging from (G). Mann-Whitney U test (N=32, 39 cells from 3 biological replicates; error bars: mean±SDs).

Importantly, the CCTF1 transcription factor was previously reported to have a cyclic expression pattern (Lundgren & Bernander, 2007), with the peak of expression also in D-phase, similar to its homolog in the related species *Saccharolobus islandicus* (Gomez-Raya-Vilanova *et al*, 2025). These results imply that the temporal change in PAN expression, mediated in part by changes in the levels of the cell cycle regulated transcription factor CCTF1, may underlie the switch in CdvB stability that occurs as cells move from a ring assembly stage (where CdvB is relatively stable) to the constriction phase (when CdvB is unstable).

## Discussion

The onset of cytokinesis in *Sulfolobus* requires rapid proteasomal-mediated degradation of the non-contractile CdvB polymer in the division ring (Tarrason Risa *et al*, 2020). Thus, proteasomal degradation plays a key role in the commitment of *Sulfolobus* cells to division. Here, using an *in vivo* flow cytometry-based degradation assay to search for sequences in CdvB about to confer cyclic degradation on a fusion protein, we mapped the regulation to the C-terminal region of CdvB. The tail of CdvB is sufficient to recapitulate the rapid degradation from D-phase to G1 phase. Importantly, this part of the protein is also absent from the other *Sulfolobus* ESCRT-III paralogs, which are degraded in G1 to S-phase of the cell cycle, once their transcription has ended. Examining individual truncation constructs revealed two important features of this C-terminal tail: (i) The C-terminal region of CdvB has a high basal protein degradation rate across all stages of the cell division cycle, implying the presence of general degron signals; (ii) the presence of the C-terminal broken winged helix domain stabilizes the protein prior to division.

These observations imply that the broken winged helix of CdvB temporarily inhibits its degradation pre-division, contributing to the observed cell cycle-dependent expression profile. While an earlier study showed that disassembly of the CdvB polymer is likely a prerequisite for its proteasome-mediated degradation (Hurtig *et al*, 2023; Liu *et al*, 2025), the stabilization due to the broken winged helix proved to be independent of CdvB polymerization, since fusion proteins that lack the ESCRT-III core domain are not associated with the division ring, but still undergo a switch in stability as they pass from pre-division into G1. Intriguingly, this is the same region of the protein that was previously shown to bind to CdvA (Samson *et al*, 2011). This suggests the possibility that, during ring assembly, which is triggered by the accumulation of CdvA in early division phase of the cell cycle, the ability of the E3B helix to complete the winged helix fold may stabilize CdvB to prevent its proteasome-mediated degradation.

Our data also point to additional factors contributing to the change in CdvB degradation rate as cells undergo division. As cells pass from late G2 into G1, there is a change in the level of PAN - the disassembly factor that unfolds proteins as a prelude to proteasome-mediated degradation. Moreover, PAN is required to prevent CdvB persisting until G1, implying that these cyclic changes in the expression of PAN play a role. We speculate that these different factors including degradation signals intrinsic to CdvB, its protection from degradation when present in a polymer, and cell cycle changes in the levels of PAN, all contribute to the switch-like change in the rate of CdvB degradation at division in *S. acidocaldarius*.

This rapid change in CdvB stability is important as it allows the initial buildup of cellular CdvB concentration during the pre-division phase as cells are constructing a template division ring, which serves as a platform for the recruitment of CdvB1 and CdvB2 (Fig. 6). As the division ring is assembled, CdvB is likely stabilized both by the presence of CdvA and by the formation of an ESCRT-III polymer, which hinders degradation by the proteasome. Subsequently, following the switch in the levels of CCTF1, Vsp4 and PAN that occur as cells progress to a state in which the DNA is segregated as a prelude to division (Parham *et al*, 2025), the CdvB polymer is then disassembled by Vps4 and separated from CdvA. This generates a pool of free CdvB monomer that can then be unfolded by PAN, which is now present at relatively high concentrations in cells, leading to the rapid degradation of the protein by the proteasome. The loss of CdvB from the division ring frees CdvB1/B2 polymers to constrict and drive cytokinesis (see Fig. 6 for the proposed model).

**Figure 6.**
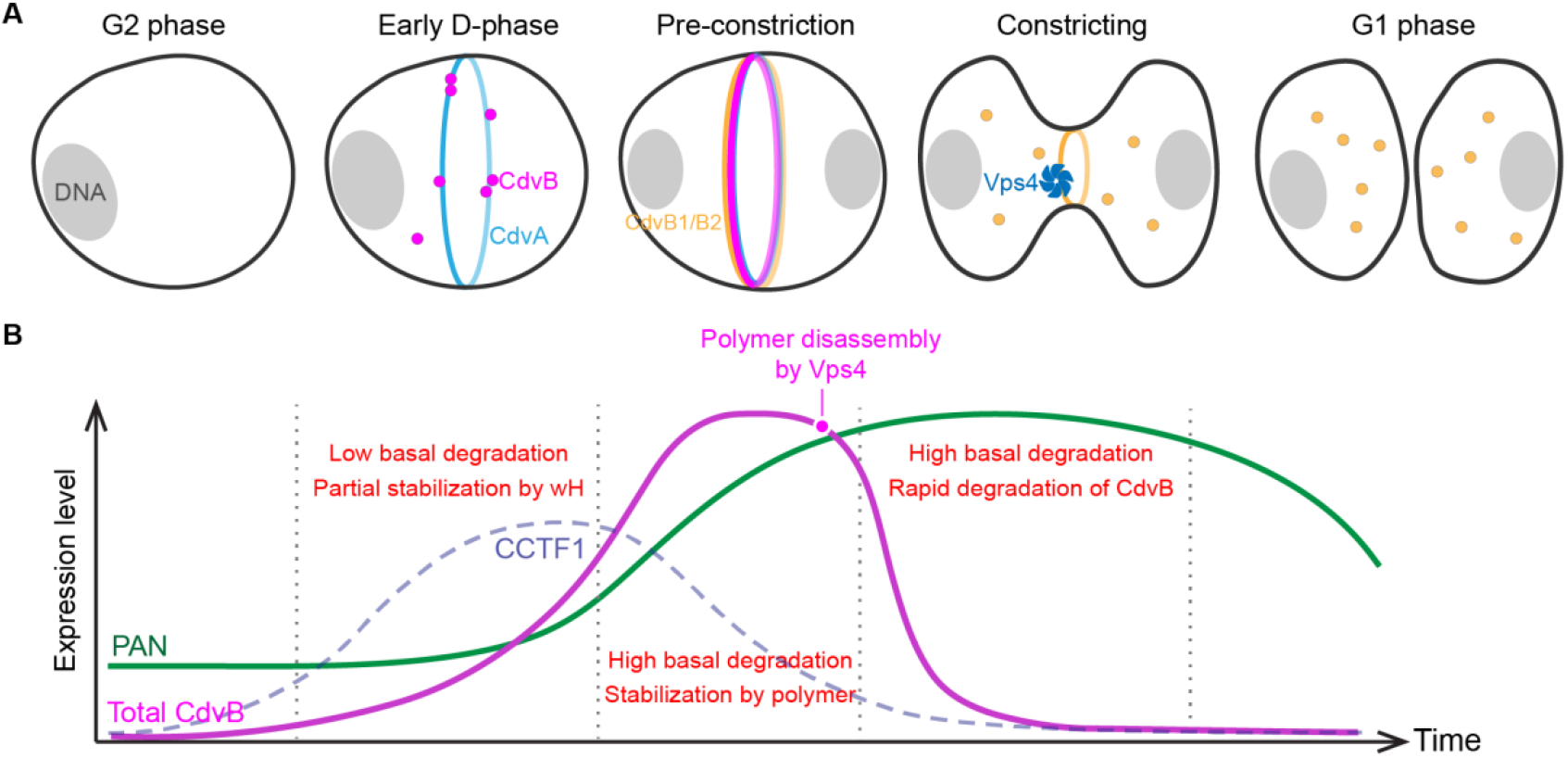
Proposed model for cyclic degradation of CdvB. **(A)** Graphic schemes showing the key stages from G2 phase to D-phase and completion of cell division. **(B)** Schematic diagram of the change in CdvB (magenta), PAN (green) and postulated CCTF1 (blue dashed line) abundance during the division phase.

Notably, a recent study in closely related species *Saccharolobus islandicus* suggested that cell cycle-dependent phosphorylation of the 20S proteasome subunits can regulate its assembly and interaction with PAN to modulate degradation activity during division (Huang *et al*, 2025). While these phosphorylation sites are not conserved across Sulfolobales, their effect is expected to mirror the cyclic expression of PAN we demonstrated here in *S. acidocaldarius*. This suggests that there may be multiple levels of control that all contribute to the cyclic degradation of substrates as cells pass through division and into G1. In this light, we also note that a residual pool of CdvB was reported to remain in *S. islandicus* cells as they divide (Liu *et al*, 2025), even though CdvB degradation is required for division in this organisms as it is in *S. acidocaldarius*. This implies subtle differences in the kinetics of CdvB degradation across lineages. Taken together, these findings highlight the evolutionary plasticity of the proteasome regulatory strategies that control the cell cycle progression.

In conclusion, in this study we demonstrate that archaea can achieve robust cyclic degradation of a key cell division machinery during each round of the cell cycle through a combination of substrate-specific molecular signals, polymer dynamics, and the cyclic expression of components of the protein degradation machinery. Our results provide a paradigm of a potential proteasome-mediated cell cycle regulation mechanism prior to the evolution of CDK/cyclins and E3 ubiquitin ligases. In conjunction with the transcriptional control of cell cycle-dependent gene expression (Lundgren & Bernander, 2007; Li *et al*, 2023; Huang *et al*, 2025), these molecular mechanisms can constitute an ordered cell cycle system that is functionally analogous to the case in eukaryotes.

## Materials and Methods

### Cell culture and growth conditions

All *S. acidocaldarius* strains were grown in Brock medium at 75 °C, pH 3.0 with constant shaking. Growth media is supplemented with 0.1% NZ-amine, 0.2% sucrose and with sulfuric acid used to adjust the pH to 3 (Brock *et al*, 1972). MW001, a uracil auxotrophic *S. acidocaldarius* strain (Wagner *et al*, 2012) used as the background strain in this study was grown with 4 mg/L uracil supplemented.

For all experiments, cells were grown to OD600 of 0.1-0.2, corresponding to the exponential growth phase. Cells were treated with 10 μM or 1 μM bortezomib (Abcam catalog number ab142123; stock solution 100 μM in DMSO), for proteasomal inhibition experiments. Induction of protein expression was performed by supplementation with an addition of 0.2% arabinose.

Synchronisation of cells was performed by supplementing media with 2 mM acetic acid for 4.5 hours. Cells were released from synchronisation by pelleting at 5000 rcf for 4 min. Cell pellets were washed by resuspending in fresh media and pelleting at 5000 rcf for 4 min three times, before being diluted back into fresh, pre-warmed Brock medium. Cells were fixed on ice by stepwise addition of ice-cold ethanol to the final concentration of 70%. Fixed samples were stored at 4 °C for up to three months.

### Molecular genetics

CdvB protein truncation fragments were procured commercially (gBlocks gene fragment, IDT). All fragments were designed with 30 bases of overlap with the plasmid backbone at both 5’ and 3’ ends. Fragments were assembled using Gibson Assembly (New England Biolabs, E5510S) in pSVAara-FX-stop plasmid double digested with NcoI and XhoI. Similarly, cloning of saci_0800-HA overexpression construct was performed by Gibson assembly with custom synthesised gBlock containing codon optimised *saci_0800* fragment flanked with 30 base pairs of overlapping sequencing for insert and double digested (with NcoI and XhoI) pSVAara-FX-HA plasmid as the receiving vector. All cloned plasmids were verified by Sanger sequencing. Plasmids were methylated *in vivo* by transforming into *E. coli* ER1821 (New England Biolabs). Methylated plasmids were then used for transforming electrocompetent MW001 cells by electroporation (2000 V, 25 μF, 600 ohms, 1 mm). Positive colonies were selected on gelrite-Brock plates without uracil supplementation, followed by validation of PCR genotyping with Sanger sequencing.

### Immunostaining

1 mL of fixed cells was pelleted at 8000 rcf for 3 min. Cell pellets were resuspended in PBS supplemented with 0.2% Tween-20 and 3% bovine serum albumin (PBSTA) and washed twice. After washing, cells were resuspended in 200 μL of primary antibodies (Supplementary Table S1) in PBSTA overnight at 23 °C and 400 rpm. Cells were washed three times with PBSTA and resuspended in 200 μL of 1:200 secondary antibodies in PBSTA and 200 μg/ml Concanavalin A conjugated to Alexa Fluor 647 for S-layer labelling (ThermoFisher, C21421) for 3h at 23 °C and 400 rpm shaking. Cells were washed three times with PBSTA and resuspended in PBSTA with 2 μM Hoechst 33342 (Invitrogen Cat, H3570) for visualisation of DNA for flow cytometry or 2 μM DAPI (Invitrogen, D3571) for microscopy imaging.

### Flow cytometry

Fixed cells were stained as described above. All flow cytometry analysis was performed on a BD Biosciences LSRFortessa. Laser wavelengths of 355 nm, 488 nm, 561 nm, and 633 nm were used together with filters 450/50 ultraviolet, 525/50 blue, 582/15 yellow-green, and 710/50 red respectively. Additionally, side scatter and forward scatter signals were recorded. Collected data were analysed using FlowJo v10.10. Single cells were first selected by excluding the off-diagonal data points in the height vs. area scatter plot in the Hoechst channel. For HA-tag intensity measurement, staining background was subtracted using the average intensity in the corresponding gates from the MW001 strain which serves as a negative staining control. The D-phase population was selected by gating the highest CdvA intensity populations with 2N DNA content which typically correspond to ∼5% of total cells; G1 cells were gated by selecting populations with 1N DNA content using the Hoechst channel.

### Confocal microscopy

Imaging was performed on a Nikon Eclipse Ti2 inverted microscope equipped with a Yokogawa SoRa scanner unit and Prime 95B scientific complementary metal-oxide semiconductor (sCMOS) camera (Photometrics). For this, cells were imaged in Lab-Tek #1 chambered coverglass coated with 1% polyethyleneimine for 2 h at 37°C. Images were acquired using 100x oil immersion objective (Apo TIRF 100X/NA=1.49, Nikon). The 2.8x magnification lens in the SoRa unit allowed a total magnification of 280x. For labelled proteins and DNA labels, cells were exposed to laser for 50 to 200 ms and 500 ms respectively. Laser power was adjusted accordingly to prevent saturation of pixels. Data for the z-axis were gathered by taking 10 frames with 0.22 μm step. Image analysis and z-axis sum or maximum projections were performed with Fiji.

### Western blotting

Cells were lysed in 1x Laemmli buffer (Biorad) followed by incubation of samples at 99°C for 10 min and centrifugation to remove cell debris. Samples were then loaded on a NuPAGE 4 to 12% Bis-Tris gels (Invitrogen) and ran at 150 V with MES running buffer. Transfer of proteins to the nitrocellulose membrane was done by 100 V for 1 hour in the cold room. Membranes were blocked with 5% milk in PBS supplemented with 0.2% Tween 20 (PBST). After blocking, membranes were incubated with primary antibodies (Supplementary table S1) in PBST at 4°C overnight. The Sulfobolus PAN antibody serum was shared by Dr. Nick Robinson (Lancaster University). Membranes were washed three times with PBST for 5 minutes after which membranes were incubated with 1:10000 secondary antibodies in PBST at room temperature for 2 hours. Signal from proteins was recorded by exposing membranes using the Bio-Rad ChemiDoc system, and the band intensity was analyzed by the gel analysis tool in Fiji.

### RNA extraction and RT-qPCR

For total RNA extraction, the cell pellet from 20 mL of culture was resuspended with 750 μL of TRIzol (Invitrogen) and the subsequent chloroform extraction and RNA precipitation were performed following the manufacturer’s protocol. The total RNA pellet was then dissolved in 100 μL of 1x TURBO DNase digestion buffer and treated with TURBO DNase (0.04 U/μL; Invitrogen) for 20 min at 37 °C. The RNA was then purified by using the RNeasy kit (Qiagen) and determined the purity and concentration with nanodrop before storage in −70 °C.

RT-qPCR reaction was prepared by using Luna One-step RT-qPCR kit (New England Biolab) with 5 to 10 ng of total RNA per reaction and ran on the ViiA 7 Real-Time qPCR system (Applied Biosystems). The relative RNA level was quantified by the 2^ΔΔCt^ method using SecY as the housekeeping gene. The primer pairs used are summarized in Supplementary table S2.

### Quantification and statistical analyses

All data processing and statistical analysis were performed using Microsoft Excel and GraphPad Prism 10 software.

## Author contribution

J.T., Y.-W.K. and B.B. conceived the project. J. T. generated all LacS fusion strains and performed the LacS fusion experiments with data analysis performed by Y.-W.K. and J.T. Proteasome inhibition and PAN mutant experiments were performed by J.T. and Y.-W.K. Y.-W.K. performed CCTF1 overexpression and PAN synchronization experiments. S.F. assisted with generation of strains. B.B. supervised the work. Y.-W.K, B.B. and S.F. wrote the manuscript with input from all authors.

## Acknowledgement

All flow cytometry experiments were performed at the Medical Research Council Laboratory of Molecular Biology Flow Cytometry facility, and we would like to thank members of the Flow Cytometry Facility for their technical support. We would also like to thank Dr. Nick Robinson (Lancaster University) for sharing the Sulfolobus PAN antibody. Y-WK was supported by an EMBO postdoctoral fellowship (ALTF 903-2021) and by the Medical Research Council - Laboratory of Molecular Biology (MC_UP_1201/27); SF was supported by the Wellcome Trust (222460/Z/21/Z); BB received support for work in Sulfolobus from the MedicalResearch Council - Laboratory of Molecular Biology (MC_UP_1201/27), the Wellcome Trust (222460/Z/21/Z), and the Life Sciences Moore-Simons Foundation (735929LPI).

